# Terminal complement complexes with or without C9 potentiate antimicrobial activity against *Neisseria gonorrhoeae*

**DOI:** 10.1101/2025.01.16.633325

**Authors:** Evan R. Lamb, Alison K. Criss

## Abstract

The complement cascade is a front-line defense against pathogens. Complement activation generates the membrane attack complex (MAC), a 10-11 nm diameter pore formed by complement proteins C5b through C8 and polymerized C9. The MAC embeds within the outer membrane of Gram-negative bacteria and displays bactericidal activity. In the absence of C9, C5b-C8 complexes can form 2-4 nm pores on membranes, but their relevance to microbial control is poorly understood. Deficiencies in terminal complement components uniquely predispose individuals to infections by pathogenic *Neisseria,* including *N. gonorrhoeae* (Gc). Increasing antibiotic resistance in Gc makes new therapeutic strategies a priority. Here, we demonstrate that MAC formed by complement activity in human serum disrupts the Gc outer and inner membranes, potentiating the activity of antimicrobials against Gc and re-sensitizing multidrug resistant Gc to antibiotics. C9-depleted serum also disrupts Gc membranes and exerts antigonococcal activity, effects that are not reported in other Gram-negative bacteria. C5b-C8 complex formation potentiates Gc sensitivity to azithromycin but not lysozyme. These findings expand our mechanistic understanding of complement lytic activity, suggest a size limitation for terminal complement-mediated enhancement of antimicrobials against Gc, and suggest complement manipulation can be used to combat drug-resistant gonorrhea.

**Importance:** The complement cascade is a front-line arm of the innate immune system against pathogens. Complement activation results in membrane attack complex (MAC) pores forming on the outer membrane of Gram-negative bacteria, resulting in bacterial death. Individuals who cannot generate MAC are specifically susceptible to infection by pathogenic *Neisseria* species including *N. gonorrhoeae* (Gc). High rates of gonorrhea and its complications like infertility, and high-frequency resistance to multiple antibiotics, make it important to identify new approaches to combat Gc. Beyond direct anti-Gc activity, we found the MAC increases the ability of antibiotics and antimicrobial proteins to kill Gc and re-sensitizes multidrug-resistant bacteria to antibiotics. The most terminal component, C9, is needed to potentiate the anti-Gc activity of lysozyme, but azithromycin activity is potentiated regardless of C9. These findings highlight the unique effects of MAC on Gc and suggest novel translational avenues to combat drug-resistant gonorrhea.

## Introduction

The complement system is a predominant arm of innate immunity that is a front-line defense for combating pathogens (1–7). Complement components are abundant in serum and found at most tissues and mucosal surfaces (8–11). Complement activation is robustly initiated by IgG and IgM binding, and the resulting catalytic cascade promotes effector functions including leukocyte activation and opsonophagocytosis of C3b-labeled targets by phagocytes (1, 7, 10, 12). Complement directly kills pathogens by forming membrane attack complex (MAC) pores in target membranes (1, 13–15).

The MAC is generated by progressive membrane insertion of the terminal complement components C5b through C8 and subsequent polymerization of C9, resulting in 10-11nm pores (13, 14, 16, 17). The 5nm C9 transmembrane domains are not predicted to span beyond the Gram-negative outer membrane to targets deeper in the bacterial cell (14, 16). However, in *Escherichia coli*, outer membrane disruption alone is insufficient to drive bacterial death by the MAC, whereas inner membrane disruption is essential (16, 18–20). Therefore, foundational biologic questions remain as to how the MAC promotes bactericidal activity. Furthermore, C5b-C8 complexes, without poly-C9, can themselves cluster in membranes, forming smaller pores (∼2-4nm) that lyse liposomes and erythrocytes, and kill nucleated cells. Effects and mechanisms of C5b-C8 complexes on Gram-negative bacteria remain to be fully investigated (21–26).

Deficiencies in the complement system result in increased susceptibility to certain infections (1, 2). In particular, deficiencies in C5 through C9 result in a 1,000- to 10,000-fold increased risk for invasive meningococcal disease by *Neisseria meningitidis* and >300-fold increased susceptibility to local and disseminated infection by *N. gonorrhoeae* (1, 27). In turn, pathogenic *Neisseria* attempt to evade complement-mediated killing by hijacking host-derived complement inhibitors C4b-binding protein, factor H, sialic acid, and vitronectin, evading antibody recognition by phase and antigenic variation, and meningococcal capsule production (1, 28–36).

*N. gonorrhoeae* (the gonococcus, Gc) causes an estimated 82-100 million cases of gonorrhea annually worldwide (37–39). Gonorrhea is an urgent public health threat due to rapidly rising case numbers along with increasing antibiotic resistance (39–42). Gc infection is characterized by mucosal inflammation, resulting in an influx of neutrophils and serum transudate (43). If left untreated, or if treatment is ineffective due to antibiotic resistance, collateral tissue damage can cause serious sequelae including pelvic inflammatory disease, ectopic pregnancy, endocarditis, and infertility (1, 43).

Gonococci have been isolated that are resistant to all classes of antibiotics that have been used for treatment, including macrolides, fluoroquinolones, tetracyclines, and β-lactams. Extensively-drug resistant Gc with lowered susceptibility to extended spectrum cephalosporins are circulating worldwide (40–42). Resistance is conferred by mutation of the antibiotic’s target, reduced uptake via mutations in the outer membrane porin, and increased efflux pump production (44–47). As in other Gram-negatives, the outer membrane is a barrier preventing access to deeper sites in the bacterial cell (45, 48–52).

MAC-mediated disruption of the outer membrane can enhance bactericidal activity of antimicrobials against Gram-negative bacteria (53–55). In this model of MAC-mediated potentiation, antimicrobials that are excluded by the outer membrane gain access to the inside of the bacterial cell by traversing through the MAC pore, similar to pharmacologic strategies of enhancing antibiotic activity by combining them with membrane-disrupting compounds (50, 56, 57). However, it is unclear whether MAC-mediated potentiation is conferred by antimicrobial transit through channels formed by the MAC pore or by generalized membrane perturbation (58). The ability of C5b-C8 complexes to potentiate antimicrobials has also not been tested.

Given these observations and the importance of complement to control *Neisseria*, we investigated how sublethal MAC deposition affected Gc susceptibility to curated antimicrobials. We demonstrate that MAC damages both the gonococcal outer and inner membranes and enhances antibiotic activity at each layer of the Gram-negative cell. Moreover, the MAC re-sensitizes a multidrug-resistant Gc strain to clinically relevant antibiotics. C9-deficient serum promotes membrane damage and antigonococcal activity of antibiotics, but does not potentiate the activity of host-derived lysozyme, implicating C5b-C8 in forming size-restricted pores in Gc. Our results reveal differences in how terminal complement restricts Gc compared with other Gram-negative bacteria and help explain how terminal complement deficiencies uniquely sensitize individuals to *Neisseria*, suggesting novel host-targeting therapeutic approaches to help combat drug-resistant gonorrhea.

## Results

### Human serum kills Gc via terminal complement component deposition

A serum bactericidal assay (SBA) was adapted to interrogate MAC disruption and antimicrobial potentiation of Gc (59). Gc was incubated with anti-lipooligosaccharide IgM, followed by addition of Ig-depleted pooled human serum as complement source; serum can contain antibodies that cross-react with Gc antigens, even in individuals with no prior Gc exposure (60). Titrating both serum and IgM concentrations resulted in significant, reproducible concentration-dependent Gc killing (Fig. 1A,B). 410ng/mL anti-Gc IgM and 2-3% serum yielded non-significant yet detectable killing (sublethal). Serum that was heat-inactivated (HI) or treated with the C5-specific inhibitor OMCI (*Ornithodoros moubata* complement inhibitor) fully lost bactericidal activity (Fig. 1A-C) (55, 61, 62). By imaging flow cytometry, C3b, C7, and C9 were on the surface of Gc incubated with IgM and active, but not HI serum (Fig. 1D-G). We conclude that Gc is susceptible to classical complement-mediated killing via the MAC in a serum- and antibody-concentration dependent manner (63).

**Figure 1.**
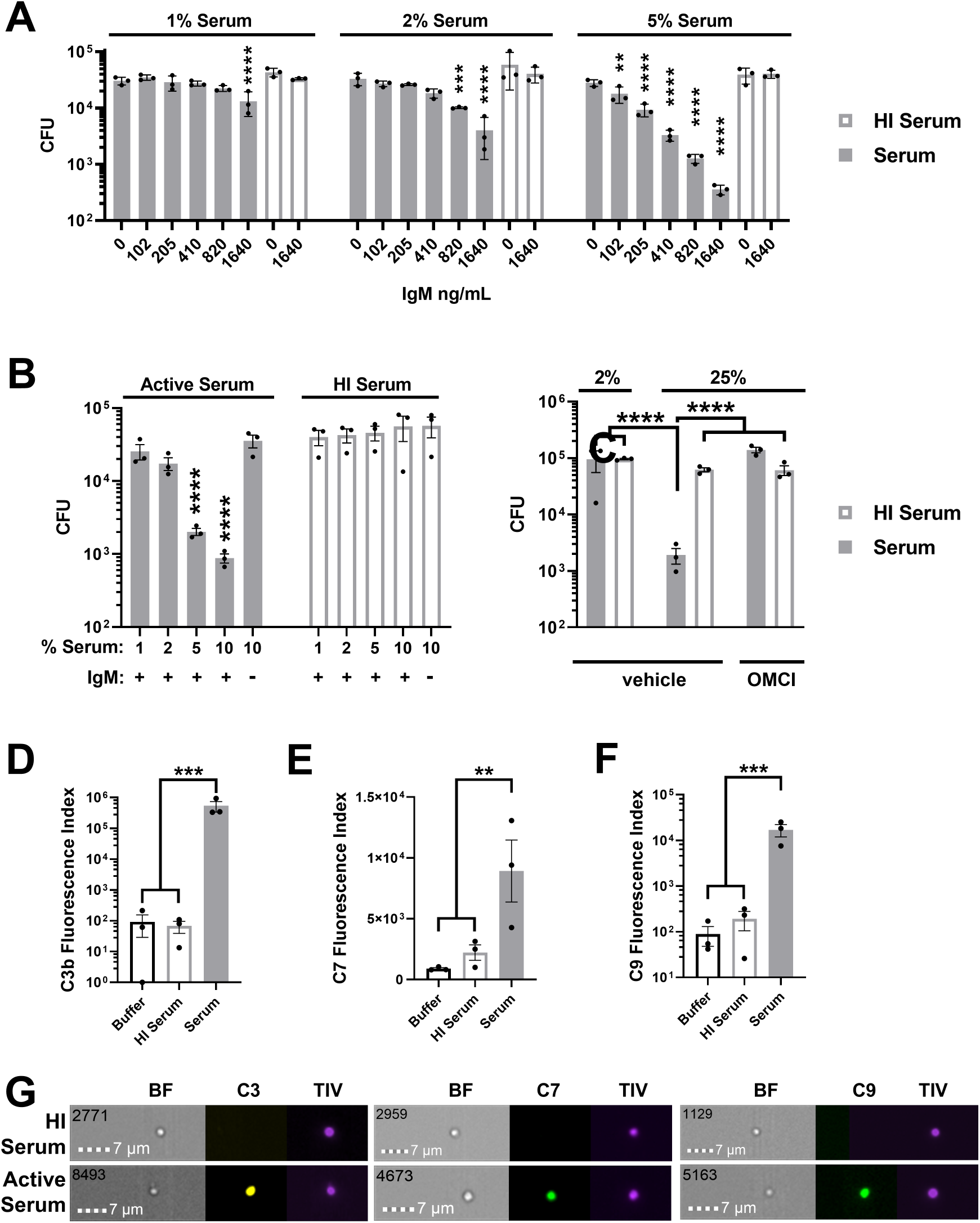
lgG/M-depleted human serum exhibits MAC-mediated bactericidal activity against Ge. **(A)** FA1090 Ge was pre-incubated with increasing concentrations of anti-Ge lgM 684, followed by incubation with active or heat-inactivated (HI) lgG/M-depleted human serum at 1, 2, or 5% final concentration. **(B)** FA1090 Ge was pre-incubated without antibody or with 41Ong/ml anti-Ge lgM, then challenged with increasing concentrations of lgG/M-depleted human serum. **(C)** FA1090 Ge was incubated with 41Ong/ml anti-Ge lgM and indicated serum concentrations with 20µg/ml of the C5 inhibitor OMCI or vehicle. In **(A-C),** CFU were enumerated from serial dilutions. **(D-G)** H041 Ge was treated with lgM for 30 min, then incubated with 2% **(D)** or 50% **(E,F)** lgG/M-depleted serum for 2hr, followed by staining and imaging flow cytometry for C3 **(D), C? (E),** or C9 **(F).** Data are presented as Fluorescence Index (median fluorescence intensity* percent positive). **(G),** representative micrographs from imaging flow cytometry of C3b, C?, and C9 binding to individual Ge. The scale bar is in the lower lefthand corner. The upper lefthand number indicates the event number of single, focused Ge out of 10,000 total events. BF= brightfield, TIV = Tag-IT Violet counterstain. Error bars are standard error of the mean. Significance was determined by 1-way ANOVA with Tukey’s multiple comparisons on log10-transformed data versus Ong/ml lgM in HI serum at indicated serum percentages **(A),** vs. 10% HI serum without lgM **(B),** or as indicated by comparison bars **(C-F).** ** = p<0.01, *** = p<0.001, **** = p<0.0001.

### The MAC disrupts both the gonococcal outer and inner membranes

The SBA conditions above were used to assess complement disruption of Gc outer and inner membranes. 1-N-phenylnapthylamine (NPN) fluoresces only upon integration into the inner membrane, following outer membrane disruption (57, 64). NPN fluorescence was significantly increased in Gc in an active complement-dependent manner (Fig. 2A). Sytox Green fluoresces upon DNA intercalation, after disruption of both outer and inner membranes (16, 55). Gc incubated with active serum, but not HI serum or buffer, showed increased Sytox fluorescence over 2hr (Fig. 2B). Endpoint Sytox Green fluorescence and area under the curve (AUC) were significantly increased in Gc exposed to active serum (Fig. 2C). We conclude that human serum damages both gonococcal outer and inner membranes.

**Figure 2.**
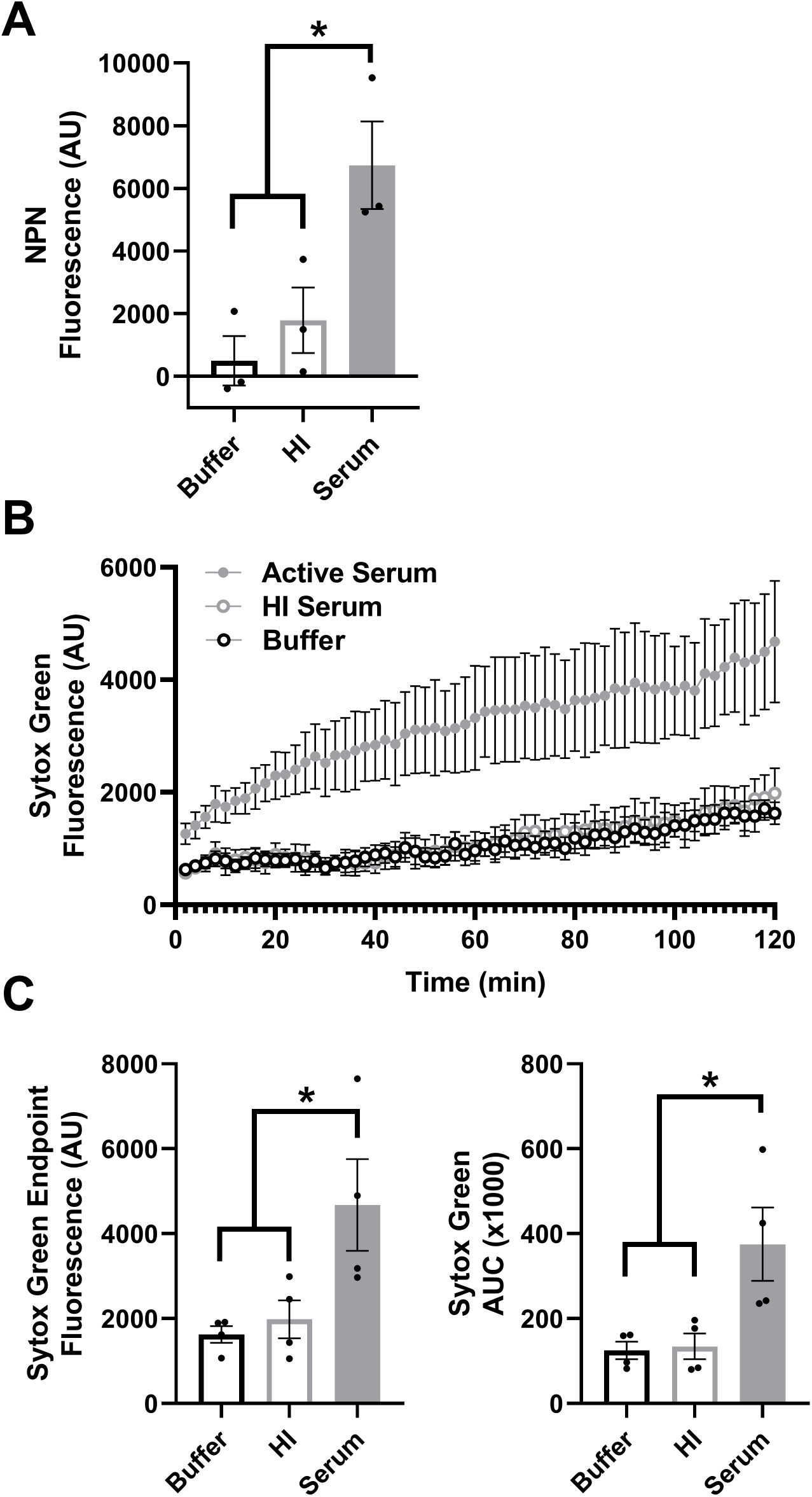
The MAC disrupts the gonococcal outer and inner membranes. (A-B) Ge was pre-incubated with anti-Ge lgM followed by incubation with active serum, heat-inactivated (HI) serum, or buffer and assessed for NPN **(A)** or Sytox Green fluorescence **(B).** NPN experiments used 1-81-S2/S-23; Sytox experiments, strain H041. **(C)** Sytox Green data from (B) displayed as fluorescence value at the end of the 2-hour incubation and calculated area under the curve (AUC) over 2 hours. Error bars are standard error of the mean. Significance was determined by 1-way ANOVA with Tukey’s multiple comparisons.*= p<0.05.

### The MAC potentiates antimicrobial activity of classically Gram-positive antibiotics

To ascertain if the MAC can enhance antimicrobial activity, we developed a modified SBA in which IgM- and serum-opsonized Gc was subsequently challenged with antibiotics or host-derived antimicrobials. As proof of concept, we assessed how MAC deposition affected the susceptibility of Gc to antibiotics that are not generally effective against Gram-negative bacteria due to poor penetration of the outer membrane coupled with active efflux: vancomycin, nisin, and linezolid (50, 53, 65).

Vancomycin targets D-Ala-D-Ala linkages of peptidoglycan. Treating FA1090 Gc with 3μg/mL vancomycin and 2% active serum reduced viability by 5,327-fold. In comparison, the viability of Gc exposed to 2% serum alone reduced by 3.7-fold; when exposed to the same concentrations of vancomycin and HI serum, viability reduced 304-fold (Fig. 3A). The statistically significant, greater-than-additive effect of active serum and antibiotic was calculated as a potentiation index, defined as the ratio of antibiotic killing in the presence of active serum versus HI serum (see Methods). A potentiation index >1.0 indicates a greater-than-additive effect from combining antibiotic and active serum. The calculated potentiation index of 3μg/mL vancomycin in 2% serum was 4.7 (Fig. 3A, Table 1). Incubation with OMCI abrogated vancomycin potentiation (potentiation index of 0.83) and was no different from incubation with HI serum, showing potentiation of vancomycin was dependent on terminal complement (Fig. 3A, Table 1). Potentiation was measured over serum and vancomycin concentrations in 2-way titration experiments (Fig. 3B). Active serum also potentiated vancomycin’s activity against the unrelated Gc strain MS11 (Supplemental Figure 1, Table 1).

**Figure 3.**
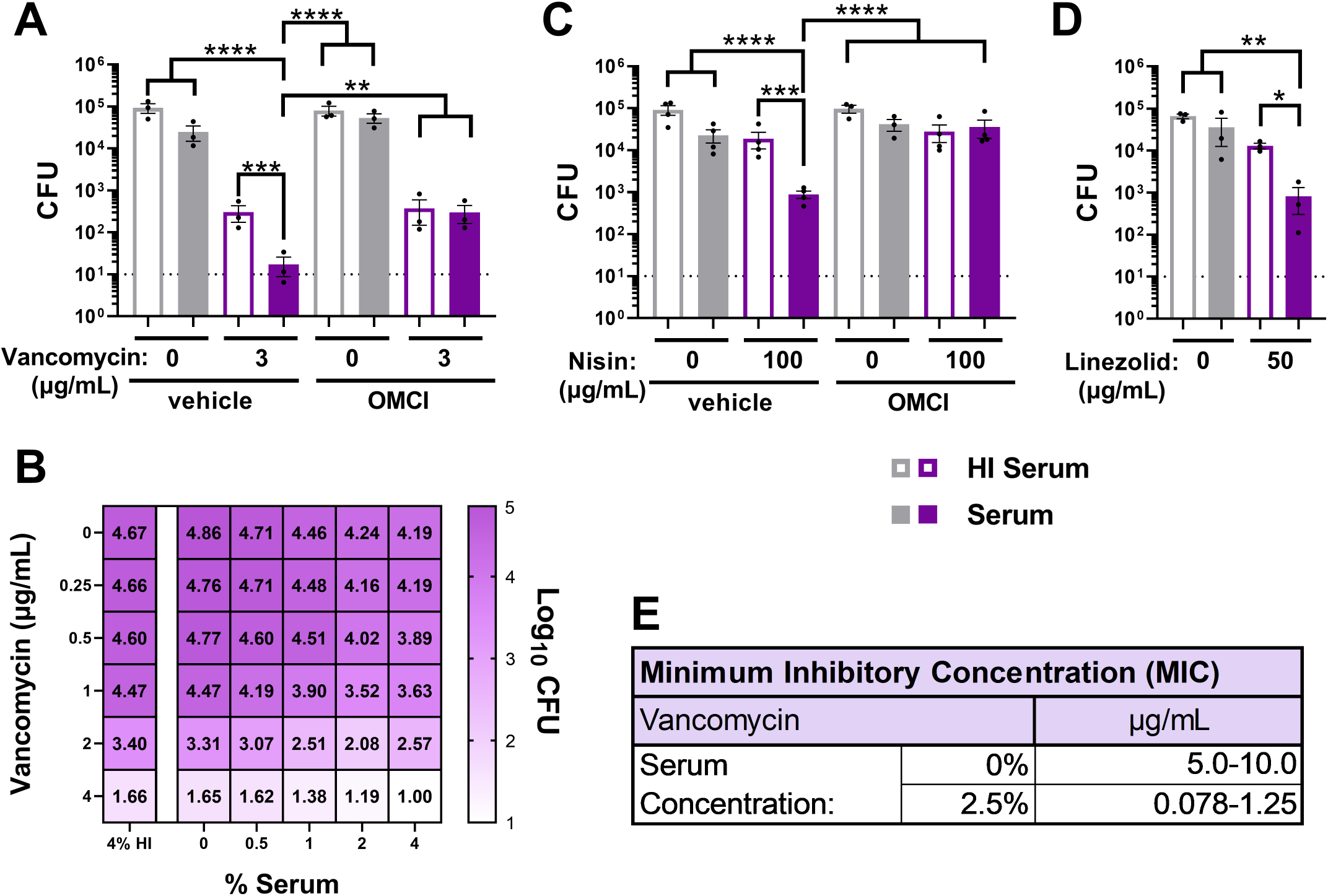
The MAC potentiates antimicrobial activity of classically Gram-positive antibiotics that act at all layers of the gonococcal cell. (A-D) FA1090 Ge was preincubated with anti-Ge lgM followed by incubation with 2% **(A,C),** 3% **(D),** or indicated concentration **(B)** of human lgG/M-depleted human serum with or without heat inactivation (HI). Ge was then incubated with the indicated antibiotic, and CFU were enumerated. Where indicated, serum was first incubated with the CS inhibitor OMCI (20µg/ml) or vehicle. Error bars are standard error of the mean. Significance was determined by 1-way ANOVA with Tukey’s multiple comparisons on Log10-transformed data. * = p<0.05, ** = p<0.01, *** = p<0.001, **** = p<0.0001. Dotted line represents minimum reportable CFUs. **(E)** FA19 Ge assayed via 16-hour minimum inhibitory concentration (MIC) broth microdilution assay over a range of vancomycin concentrations in GCBL alone or supplemented with 2.5% lgG/M-depleted human serum.

**Table 1.** Potentiation Indexes.

| **Antimicrobial (µg/mL)** | **% IgG/M-depl Serum** | **Potentiation Index** | **Potentiation Index with OMCI** | **Gc Strain** |
| --- | --- | --- | --- | --- |
| Vancomycin (3) | 2 | 4.7 | 0.83 | FA1090 |
| Nisin (100) | 2 | 3.7 | 0.33 | FA1090 |
| Linezolid (50) | 3 | 8.6 | - | FA1090 |
| Vancomycin (4) | 2 | 3.6 | - | MS11 |
| Azithromycin (4) | 2 | 174.5 | 0.90 | H041 |
| Ceftriaxone (4) | 2 | 12.5 | 0.68 | H041 |
| Zoliflodacin (0.125) | 2 | 9.7 | - | H041 |
| Doxycycline (4) | 2 | 99.8 | - | H041 |
| Gentamicin (10) | 2 | 4.8 | - | H041 |
| Lysozyme (1,000) | 2 | 3.8 | 1.0 | FA1090 |
|  | **% C9-reconst Serum** |  |  |  |
| Azithromycin (4) | 1 | 535.1 | - | H041 |
| Lysozyme (1,000) | 1 | 9.3 | - | H041 |
|  | **% C9-depl Serum** |  |  |  |
| Azithromycin (4) | 1 | 120.7 | - | H041 |
| Lysozyme (1,000) | 1 | 0.95 | - | H041 |

Given that serum causes Gc inner membrane damage (Fig. 2B,C), we next tested classically-Gram-positive antibiotics that target either lipid II in the inner membrane (nisin) or ribosomes in the cytoplasm (linezolid). In HI serum, nisin and linezolid had minimal effect on Gc viability at 100 and 50 μg/mL, respectively (Fig. 3C,D). The antigonococcal activities of nisin and linezolid were significantly increased with active serum (Fig. 3C,D) and reduced to HI serum levels when OMCI was added, indicating MAC dependence (Fig. 3C). Potentiation indexes for nisin and linezolid were 3.7 and 8.6, respectively (Table 1).

Vancomycin potentiation was independently measured using overnight broth microdilution assays for MIC determination. Addition of 2.5% active serum reduced the MIC from 5-10μg/mL to 0.078-0.123μg/mL, a 40- to 128-fold decrease (Fig. 3E). MIC broth microdilution experiments similarly demonstrated that active serum potentiated nisin activity (Supplemental Figure 2). Taken together, these data show that the MAC potentiates the activity of antibiotics that otherwise have limited activity against Gc. The use of three antibiotics with different targets and mechanisms of action emphasizes that the MAC can enable antibiotic access to all topological layers of the Gc cell.

### The MAC enhances frontline and novel antibiotic activity against multidrug-resistant Gc

Frontline antibiotic regimens for gonorrhea are ceftriaxone alone or with azithromycin, depending on local recommendations, yet resistance to these and other antibiotics is increasing (40). We determined if MAC-mediated potentiation can restore sensitivity of multidrug-resistant Gc to antibiotics using strain H041, the first isolate reported with elevated ceftriaxone resistance. H041 displays decreased susceptibility to other antibiotics, including azithromycin (45, 66). H041 Gc exposed to 2% active serum and 4μg/mL azithromycin had a 1,295-fold decrease in viability (Fig. 4A). This was a statistically significant enhancement over the effect of azithromycin alone (2% HI serum, 7.4-fold viability decrease) or when OMCI was added (Fig. 4A), resulting in a potentiation index of 174.5 (Table 1). By two-way titration, potentiation occurred over a range of azithromycin and serum concentrations (Fig. 4B). Ceftriaxone at 4μg/mL was significantly more potent at Gc killing in 2% active serum compared to HI serum, with a potentiation index of 12.5; potentiation was abrogated with OMCI (Fig. 4C).

**Figure 4.**
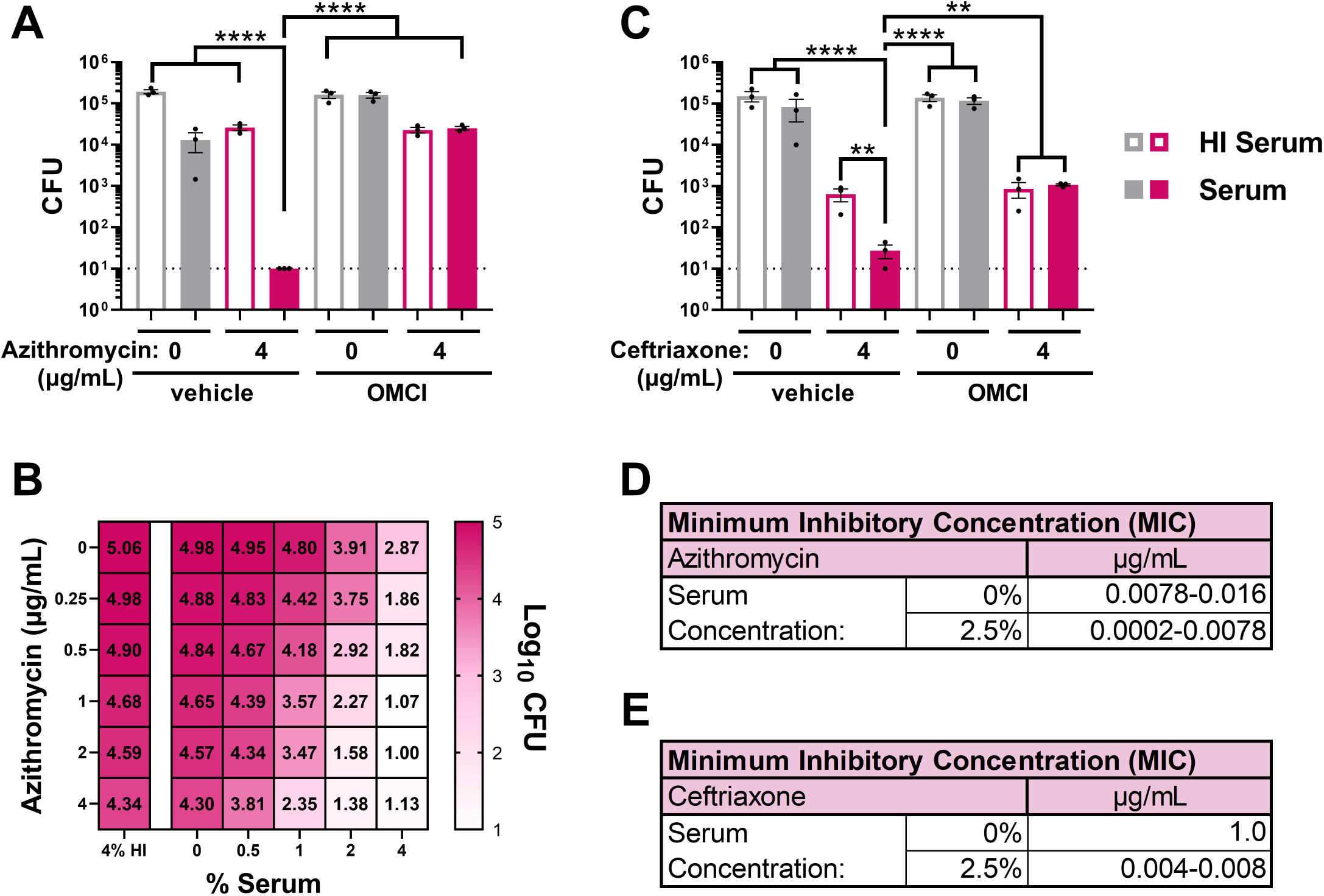
MAC-dependent increase in sensitivity and susceptibility of multidrug resistant Ge to frontline antibiotics. (A-C) H041 Ge was pre-incubated with anti-Ge lgM followed by incubation with 2% **(A,C)** or indicated concentration (**8**) of lgG/M-depleted human serum, with or without heat-inactivation (HI). Ge was then incubated with the indicated antibiotic, and CFU were enumerated. Where indicated, serum was first incubated with the CS inhibitor OMCI (20µg/mL) or vehicle. Error bars are standard error of the mean. Significance was determined by 1-way ANOVA with Tukey’s multiple comparisons on Log_10_-transformed data. ** = p<0.01, **** = p<0.0001. Dotted line represents minimum reportable CFUs. **(D,E)** FA19 Ge **(D)** or H041 Ge **(E)** were assayed via 16-hour minimum inhibitory concentration (MIC) broth microdilution assays over a range of azithromycin or ceftriaxone concentrations in GCBL alone, or supplemented with 2.5% lgG/M-depleted human serum.

By broth microdilution, the average MIC for azithromycin dropped in the presence of 2.5% active serum by 22-fold (0.0078-0.016μg/mL without serum vs. 0.0002-0.0078μg/mL with serum) (Fig 4D). Adding serum decreased the ceftriaxone MIC by 125-250-fold, from 1μg/mL to 0.004-0.008μg/mL, which is below the 0.25μg/mL susceptibility breakpoint for Gc (Fig. 4E) (67). Serum also potentiated ceftriaxone against multiple resistant Gc strains (Supplemental Figure 2). We conclude that MAC deposition renders Gc more sensitive to clinically relevant antibiotics, reducing MICs below clinically relevant breakpoints for drug-resistant strains (67, 68).

The first-in-class antibiotic zoliflodacin, a DNA gyrase inhibitor, is a promising new therapeutic for gonorrhea (ClinicalTrials.gov ID NCT03959527) (69). 2% serum significantly enhanced the activity of 0.125μg/mL zoliflodacin against H041 Gc, with a potentiation index of 9.7 (Fig. 5A, Table 1). Serum also potentiated the activity of doxycycline, currently recommended for post-exposure prophylaxis by the CDC despite a high frequency of circulating resistance in Gc (70–72), and gentamicin, currently recommended for uncomplicated urogenital infection with ceftriaxone-resistant Gc or in patients with cephalosporin sensitivity (73–75). For H041 Gc with 2% serum compared with HI serum, 4μg/mL doxycycline reduced bacterial viability 317-fold with a potentiation index of 99.8, and 10μg/mL gentamicin reduced viability 1,656-fold with a potentiation index of 4.8 (Figs. 5B,C; Table 1). Thus, new antibiotics and antibiotic treatment regimens for gonorrhea can be potentiated with human serum.

**Figure 5.**
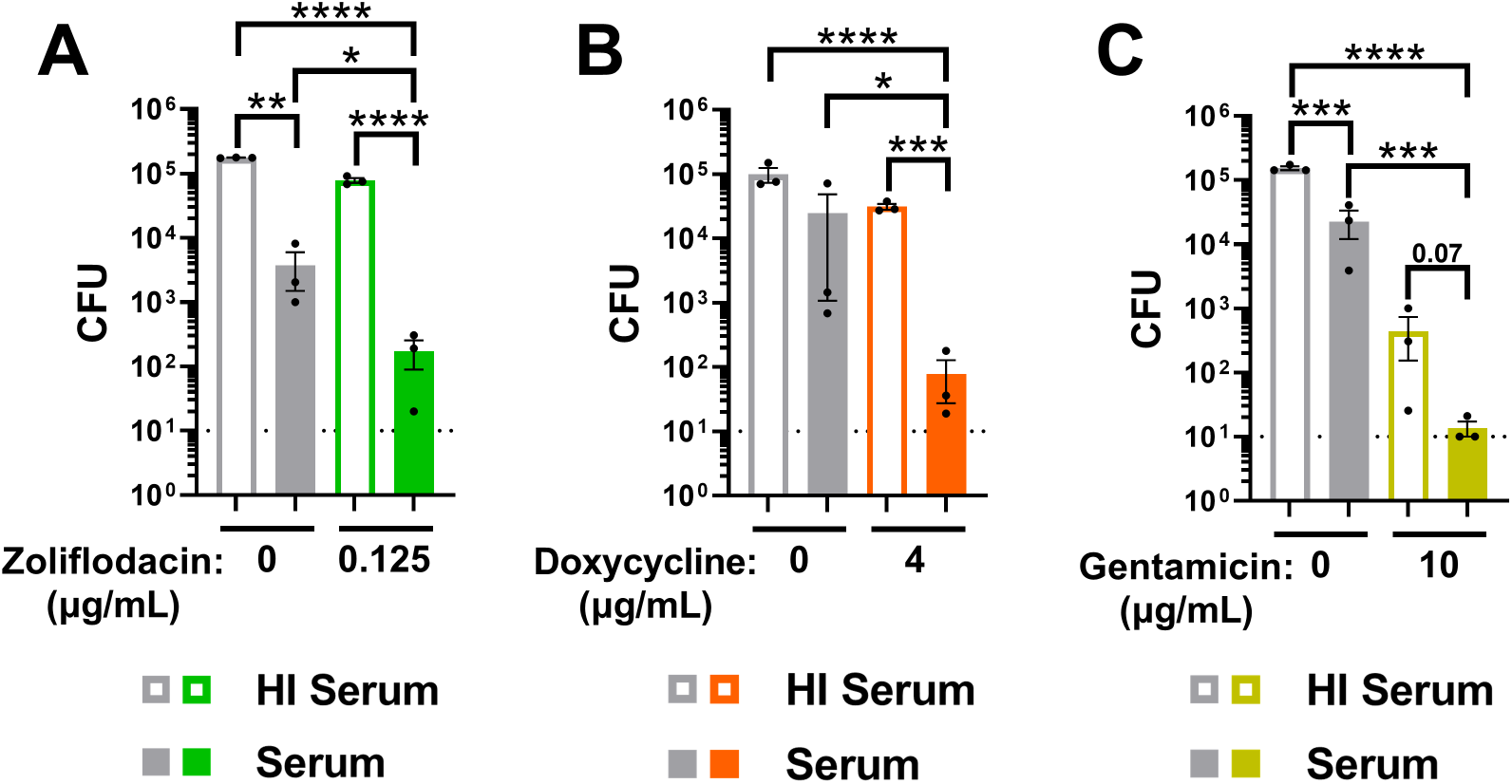
The MAC enhances the antigonococcal activity of new antibiotics and antibiotic regimens. H041 Ge was preincubated with anti-Ge lgM followed by incubation with 2% lgG/M-depleted human serum with or without heat-inactivation (HI). Ge was then incubated with zoliflodacin **(A),** doxycycline **(B),** or gentamicin **(C),** followed by CFU enumeration. Error bars are standard error of the mean. Significance was determined by 1-way ANOVA with Tukey’s multiple comparisons on Log_10_-transformed data. * = p<0.05, ** = p<0.01, *** = p<0.001, **** = p<0.0001. Dotted line represents minimum reportable CFUs.

### C5b-C8 complement complexes promote measurable bactericidal activity and damage the gonococcal outer and inner membranes

C5b-C8 complexes have been reported to form ∼2-4nm diameter pores in liposomes and eukaryotic membranes (21, 23, 24). Without C9, these smaller complexes are expected to interact differently with target membranes due to fewer transmembrane domains and their smaller size (16, 19, 24). In *E. coli*, serum depleted of C9 results in diminished outer membrane damage and little to no measurable bactericidal activity or inner membrane damage, compared to C9-replete serum (19, 22). In contrast, the viability of H041 Gc exposed to IgM and 25% C9-depleted serum was decreased by 1.2 logs compared with HI serum (Fig. 6A). Although serum reconstitution with C9 to native levels (60μg/mL) further enhanced bactericidal activity (4.1 log decrease in viability; Fig. 6A) (25), these results demonstrate that serum without C9 retains direct antigonococcal activity.

**Figure 6.**
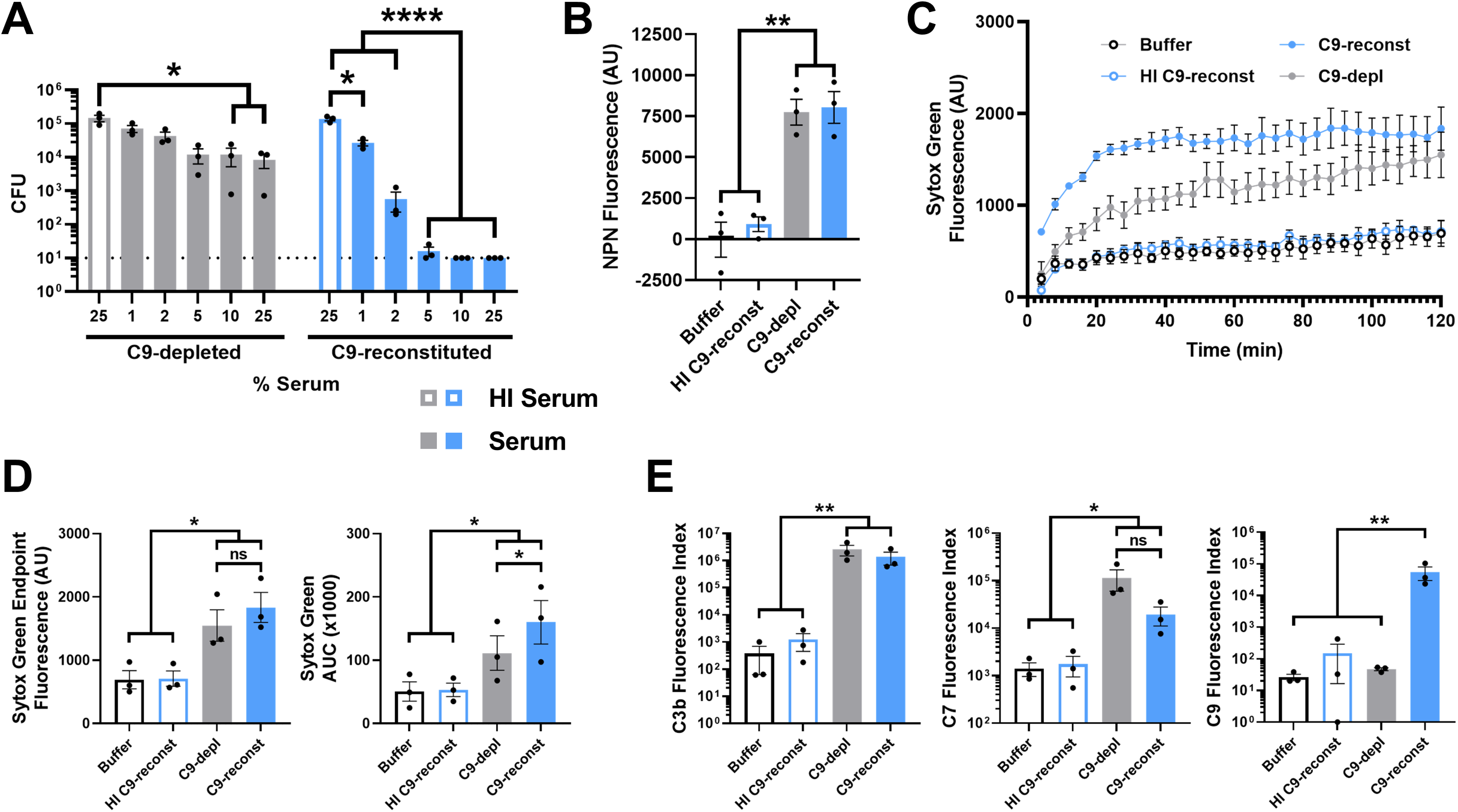
C5b-C8 complement complexes promote measurable antigonococcal activity and damage the Ge outer and inner membranes. **(A)** H041 Ge was preincubated with anti-Ge lgM, followed by incubation with the indicated concentration of C9-depleted or C9-reconstituted serum with or without heat-inactivation (HI), and CFU were enumerated. Dotted line represents CFU limit of detection. **(B-C)** Ge was pre-incubated with lgM followed by incubation with buffer, C9-depleted human serum, or C9-reconstituted human serum with or without heat inactivation. NPN **(B)** or Sytox Green fluorescence **(C)** was measured as in Figure 2. NPN experiments used FA1090/S-23, while Sytox experiments used H041. **(D)** Sytox Green data from **(C),** displayed as fluorescence value at the end of the 2hr incubation and as area under the curve (AUC) over 2hr. **(E)** H041 Ge was treated with lgM for 30min, then incubated with 2% (for C3b) or 50% (for C? and C9) lgG/M-depleted serum for 2hr. Imaging flow cytometry for the indicated complement component was conducted as in Figure 1. Data are presented as Fluorescence Index (median fluorescence intensity* percent positive). Error bars are standard error of the mean. Significance was determined by 1-way ANOVA with Tukey’s multiple comparisons on Log10-transformed data **(A,D,E)** or as 1-way ANOVA with Tukey’s multiple comparisons **(B).** * = p<0.05, ** = p>0.01, *** = p<0.001, **** = p<0.0001, ns = not significant.

To uncover how C5b-C8 complexes affect Gc outer and inner membranes, we measured NPN and Sytox Green fluorescence, respectively, as in Figure 2 (16, 55, 57, 64). Gc incubated with 50% C9-depleted or C9-reconstituted serum were indistinguishable in NPN fluorescence, and both were significantly greater than Gc in HI serum or buffer (Fig. 6B). Sytox Green fluorescence increased over 2 hours following incubation with 2% C9-depleted or C9-reconstituted serum (Fig. 6C). Endpoint Sytox Green fluorescence was not significant between C9-depleted and C9-reconstituted sera. However, there was a significant increase in Sytox Green AUC for Gc incubated with C9-reconstituted serum compared to C9-depleted serum (Fig. 6C,D). Endpoint and AUC intensities were significantly lower for Gc incubated in buffer or with HI C9-reconstituted serum compared to active C9-depleted or C9-reconstituted serum (Fig. 6D). Using flow cytometry on single bacteria (59), we confirmed that Gc exposed to C9-depleted and C9-reconstituted serum had equivalent amounts of C3b and C7 on their surface, and both were significantly greater than buffer or HI serum controls (Fig. 6E). As expected, the C9 signal on Gc exposed to active C9-reconstituted serum was significantly higher than bacteria exposed to C9-depleted serum, HI C9-reconstituted serum, or buffer, all of which were at background levels (Fig. 6E). Thus C9-depleted serum is equivalent to C9-reconstituted serum for deposition of early (C3b) and precursor terminal (C7) complement components, and reconstitution with purified C9 allows C9 deposition into the Gc outer membrane.

Taken together, these results indicate that C5b-C8 complement complexes are sufficient to disrupt the gonococcal cell envelope and promote bactericidal activity, but C9 incorporation enhances inner membrane disruption and consequent Gc killing.

### Complement C5b-C8 complexes and full C5b-C9 MACs differentially potentiate antimicrobial activities

Given that C5b-C8 and C5b-C9 complexes both displayed antigonococcal activity, we evaluated how the presence or absence of C9 potentiated the activity of antimicrobials. Using the SBA protocol from Figure 4, H041 Gc was challenged with 1% C9-reconstituted or C9-depleted active serum or HI serum controls, followed by 4μg/mL azithromycin or vehicle. Azithromycin is a 749Da antibiotic with an estimated diameter of <2nm (76). Both C9-depleted and C9-reconstituted sera potentiated azithromycin activity against H041 Gc, with potentiation indexes of 535.1 and 120.7, respectively (Fig. 7A, Table 1).

**Figure 7.**
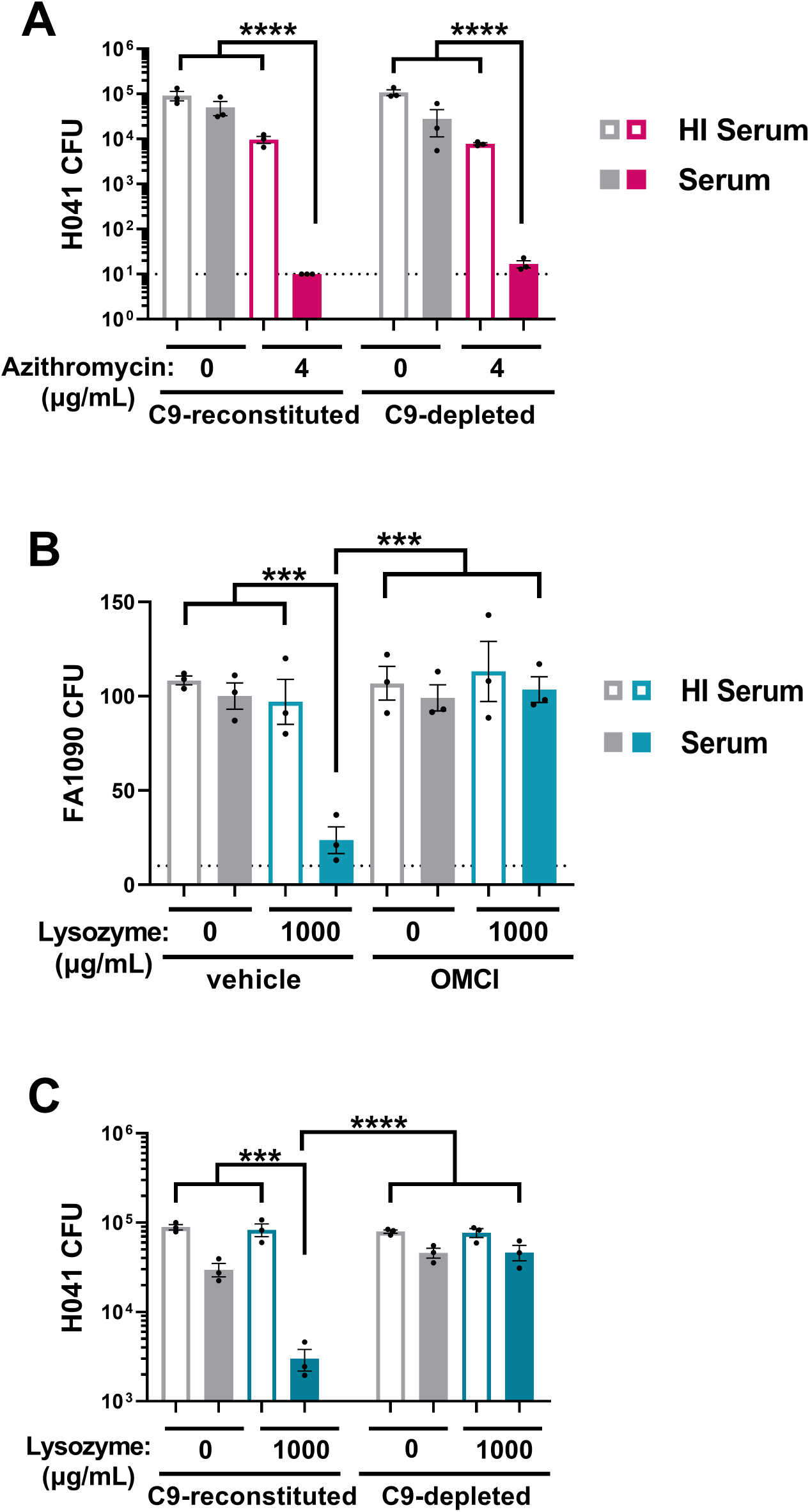
Complement C5b-C8 complexes and full C5b-C9 MAC differentially potentiate the activities of antimicrobials against Ge. H041 **(A,C)** or FA1090 **(B)** Ge was pre-incubated with anti-Ge lgM followed by incubation with 1% **(A,C)** or 2% **(B)** C9-depleted or C9-reconstituted human serum with or without heat-inactivation (HI). Ge was then incubated with azithromycin **(A)** or human lysozyme **(B,C)** and then plated for CFU enumeration. Where indicated, serum was first incubated with the C5 inhibitor OMCI (20µg/ml) or vehicle alone. Error bars are standard error of the mean. Significance was determined by 1-way ANOVA with Tukey’s multiple comparisons on Log_10_-transformed data. *** = p<0.001, **** = p<0.0001. Dotted line represents CFU limit of detection.

The peptidoglycan-degrading enzyme lysozyme has potent activity against Gram-positive bacteria with exposed cell walls, but low activity against Gram-negatives due to the outer membrane barrier (14, 48, 49, 54, 77, 78). Human lysozyme has a molecular weight of 14,300Da and a maximum diameter of ∼9nm by X-ray crystallography (79, 80). 2% active serum natively containing C9 enhanced the activity of 1000μg/mL lysozyme against FA1090 Gc with a potentiation index of 3.8; OMCI treatment abrogated the potentiation, indicating MAC dependence (Fig. 7B, Table 1). H041 Gc was resistant to killing by 1000μg/mL lysozyme when HI serum was used (Fig. 7C). Adding 1% C9-reconstituted human serum reduced Gc viability 29.7-fold, with a potentiation index of 9.3 (Fig. 7C, Table 1). In contrast, C9-depleted serum showed no potentiation of lysozyme (index of 0.95) (Fig. 7C, Table 1). We conclude that C5b-C8 and C5b-C9 complement complexes can permit small molecules such as antibiotics to bypass the Gc outer membrane, but larger molecules or antimicrobial enzymes require full C9-containing MAC pores for intracellular access.

## Discussion

Deficiencies in terminal complement components which comprise the MAC are highly predisposing to serious infections by Gc and *N. meningitidis* (1, 27). The capacity for MAC to damage neisserial membranes and enhance antimicrobial activity represents a promising avenue for combating these pathogens. Here, using laboratory and multidrug-resistant strains of Gc, we found the MAC disrupted both outer and inner membrane integrity. Beyond direct bactericidal activity, MAC enhanced the antigonococcal activity of antibiotics and rendered multidrug-resistant Gc susceptible to frontline and new antibiotic programs. Intriguingly, C5b-C8 complexes also disrupted Gc outer and inner membranes and exerted bactericidal activity. C5b-C8 complexes potentiated the activity of azithromycin, but C9 addition was necessary to potentiate lysozyme. We conclude that terminal complement components, both MAC and C5b-C8 complexes, are both directly bactericidal for Gc and also potentiate the activity of diverse antimicrobials.

As a mucosal pathogen, Gc encounters complement via serum transudate and local production by resident epithelial cells, fibroblasts, and immune cells (8, 11, 43, 81). Here, we showed that serum exposure enhances killing of Gc by antimicrobials targeting the periplasm (vancomycin, ceftriaxone, lysozyme), inner membrane (nisin), and cytoplasm (linezolid, azithromycin, zoliflodacin, doxycycline, gentamicin), which is abrogated by heat inactivation or OMCI. Thus, antimicrobial potentiation is MAC-dependent and broadly applicable to different treatment options. As C5b-C8/C9 complexes disrupt both outer and inner membranes, we conclude that terminal complement perturbs the Gc envelope to enhance antimicrobial penetration. MAC-mediated potentiation underscores the promise of membrane-disrupting therapies as adjuvants to enhance antibiotic efficacy against multidrug-resistant bacteria like Gc.

Although the MAC cannot extend past the outer membrane, inner membrane disruption is required for MAC to kill Gram-negative bacteria (16, 18–20). The exact mechanism of MAC killing remains undefined but could include generalized osmotic instability, leakage of vital intracellular factors, influx of toxic factors, homeostatic disturbance (diminished proton motive force [PMF]), and triggering of stress responses leading to bacterial death (16, 82). Several non-exclusive hypotheses can explain how MAC potentiates antimicrobial activity in Gc. First, outer membrane disruption increases the periplasmic concentration of antibiotics, which then access the cytoplasm. This possibility is supported by MAC restoring antibiotic sensitivity to multidrug-resistant Gc like H041 with more restrictive porin (44, 45). Relatedly, inner membrane disruption via MAC would also enhance cytoplasmic access of antimicrobials. Finally, inner membrane perturbation would inhibit efflux pumps that directly or indirectly require the PMF (47). Although efflux pumps are frequently upregulated in multidrug-resistant Gc (44–46), terminal complement activity would overcome their activity. Future studies can test among these hypotheses by tracking antimicrobial access to subcellular compartments.

We found that serum containing C9 was bactericidal for Gc and that C9-containing MAC disrupted Gc outer and inner membranes. Notably, C5b-C8 complexes also promoted antigonococcal activity, though less robustly. The Gc outer membrane was damaged similarly by C9-depleted and C9-reconstituted serum, while inner membrane damage by C9-depleted serum was delayed but reached the same endpoint as with C9. These findings contrast with results from *E. coli,* where C5b-C8 complexes minimally affected inner membrane integrity or bactericidal activity compared to MAC (19, 22). The uniqueness of *Neisseria* cell wall composition and integrity versus other Gram-negative bacteria may underlie these C9-dependent differences. The outer leaflet of the neisserial outer membrane is composed of lipooligosaccharide, not lipopolysaccharide (83, 84). Unlike other Gram-negative bacteria, Gc lipid membranes contain significant levels of phosphatidylcholine and differ in other phospholipid species composition (85–87). Gc lacks Braun’s lipoprotein (88) or full-length OmpA or Pal homologues, which link the outer membrane to the cell wall (89, 90). The Rcs system that senses outer membrane stress is also absent in Gc (91–93). Because Gc subverts both human cellular and humoral immunity, including resistance to neutrophils (77, 94–96), prevention of protective T_H_1 responses (97, 98), induction of B cell death and impaired antibody production (99), and phase and antigenic variation to evade antibody recognition (34, 100), complement may be the most effective arm of immunity to control Gc, and its absence greatly increases susceptibility to infection. Our findings with C9 align with epidemiologic evidence that C9 deficiencies more modestly predispose individuals to *Neisseria* compared to other terminal complement deficiencies (1, 25, 101). Beyond genetic C9 deficiencies, reduced C9 on the Gc surface could occur by bacterial recruitment of the C9 inhibitor vitronectin (1, 26, 32, 33, 102).

If terminal complement pores directly enable intracellular access to bacteria, then 10-11nm MAC pores would allow access of some antimicrobials that would be excluded by 2-4nm C5b-C8 complexes based on the antimicrobials’ diameter. We found that lysozyme was only potentiated by C9-reconstituted serum, but azithromycin was potentiated in a C9-independent manner. Thus, our results support a model in which potentiation in Gc occurs through direct transit, and that C5b-C8 complexes and MAC differentially potentiate antimicrobials in a size-dependent manner. However, the possibility remains that generalized outer membrane perturbation or ‘fracturing’ allows compounds to gain intracellular access without transiting directly through pores formed by terminal complement (58).

Our results emphasize how complement envelope perturbation could enhance anti-Gc therapeutics, including vaccines. This study used an anti-lipooligosaccharide IgM as proof of concept to drive classical complement activation on Gc (11, 59, 103). Antibody-eliciting vaccines and passive immunization with monoclonal antibodies have shown preclinical promise in preventing Gc infection in animal models and epidemiological studies (104–108). However, antibodies as immune correlates for protection have not yet been established (59, 109, 110). Even if antibodies do not drive strong bactericidal activity, our findings show that sublethal terminal complement deposition potentiates antibiotic activity. Aligning with our results, a chimeric IgM-C4b binding protein fusion increases direct killing of Gc and enhances killing by azithromycin and ciprofloxacin (55, 111). Beyond antibiotics, the finding that MAC renders Gc susceptible to killing by human lysozyme suggests that enhancing terminal complement deposition on Gc would enhance killing of Gc at mucosal surfaces and within immune cell phagosomes where these antimicrobials are found. Antibodies and complement would work together against Gc in three ways: direct lysis, opsonophagocytic killing, and potentiating antimicrobial sensitivity within and outside cells (112).

This study emphasizes that complement-mediated control of Gc can be accomplished though both MAC and C5b-C8 complexes that potentiate existing and novel antibiotic regimens and enhance host-derived antimicrobial activity. New therapeutic approaches that exploit terminal complement are promising countermeasures to combat antibiotic-resistant gonorrhea.

## Materials and Methods

### Sex as a biological variable

Human serum was pooled from both sexes.

### Neisseria gonorrhoeae

The following Gc strains were used for this study (59): FA1090 (1-81-S2, and 1-81-S2/S-23) (113), H041 (WHO X) (44, 45), MS11 (114), and FA19 (115). The 1-81-S2 strain of Gc is an FA1090 derivative with a defined pilin antigen (116–118); S-23 is a 1-81-S2 derivative where all *opa* genes were deleted and containing a loop 6 *porB* mutation that abrogates binding of C4b-binding protein to enhance serum sensitivity (119, 120). Gc was routinely streaked on gonococcal base medium (BD Difco) plus Kellogg’s supplement I and 1.25μM Fe(NO_3_)_3_ [gonococcal base (GCB)] plates for single colonies for 14-16 h at 37°C, 5% CO_2_ (59, 121). When indicated, Gc was inoculated into GCB liquid media (GCBL) or Hanks’ Balanced Salt solution with 2% bovine serum albumin (HBSS + 2% BSA).

### Human serum complement sources

IgG/IgM-depleted pooled human serum (IgG/M-depleted serum, Pel-Freez, Catalog #34010, Lot #28341) was used as the complement source for SBA assays with native C9, flow cytometry assays, and MIC assays. IgG/M-depleted serum from lots #28341 and #15443 were used for membrane integrity assays. Use of IgG/M-depleted serum removes the potential for variable bactericidal activity conferred by different individuals’ serum (60, 122). SBA assays evaluating C9 used C9-depleted human serum (Complement Technology, Catalog #A326, Lot #10a), reconstituted to physiological concentration with 60μg/mL C9 protein (Complement Technology, Catalog # A126, Lot #13) (123). Sera were stored at −80°C until thawed on the day of use, then diluted in HBSS + 2% BSA. Sera were heat-inactivated by incubation at 56°C for 30min (61).

### Antibodies and antimicrobials

See Table 2. Antimicrobial concentrations were determined experimentally, contextualized by *in vivo* concentrations or as antibiotic breakpoints where applicable (124–130).

**Table 2.** Reagents Used in This Study.

| **Antibodies** |  |  |  |  |  |  |
| --- | --- | --- | --- | --- | --- | --- |
| **Target** | **Source** | **Catalog Number** | **Clone** | **Conjugate** | **Lot** | **Stock Concentration** |
| LOS | Sanjay Ram* | - | 6B4 | - | - | 330 µg/mL |
| (i)C3b | BioLegend | 846103 | 3E7/<br>C3b | PE | B362314 | 100 µg/mL |
| C7 | Invitrogen | MA5-34943 | 15D1 | - | ZF4349897A | 2 mg/mL |
| C9 | Novus | NBP-21612F | 22 | FITC | D162593 | 1.35 mg/mL |
| mouse IgG<sub>1-3</sub> | Jackson Immuno Research | 115-545-164 | Poly-clonal | AF488 | 152191 | 700 µg/mL |
| **Antimicrobials** |  |  |  |  |  |  |
| **Name** | **Source** | **Catalog Number** |  |  |  |  |
| Vancomycin | Caisson | V007-1GM |  |  |  |  |
| Nisin | Cayman | 16532 |  |  |  |  |
| Linezolid | Cayman | 15012 |  |  |  |  |
| Ceftriaxone | Cayman | 18866 |  |  |  |  |
| Azithromycin | Cayman | 15004 |  |  |  |  |
| Zoliflodacin | TargetMol | 1620458-09-4 |  |  |  |  |
| Doxycycline | Sigma Aldrich | D9891-1G |  |  |  |  |
| Gentamicin | Sigma Aldrich | G3632-250MG |  |  |  |  |
| Human Lysozyme | Sigma Aldrich | L1667-1G |  |  |  |  |

\* 6B4 was generated from murine hybridoma and purified by thiophilic chromatography. RRID:
AB\_2617193.

### SBAs and antimicrobial potentiation assays

Single Gc colonies were swabbed from GCB plates into GCBL, diluted to OD_550_ nm of 0.07, then diluted 2.5-fold into HBSS + 2% BSA (buffer, ∼1.8e7 CFU/mL). Bacteria (20μL) were added to 20μL of 410ng/mL 6B4 IgM in buffer in a V-bottom 96-well plate and incubated at 37°C, 5% CO_2_ for 15min. Gc-antibody mixtures were then incubated with 40μL of buffer or indicated final percentages of serum for 45min. For SBA assays without antimicrobial challenge, bacteria were mixed with 80μL of PBS for indicated times. For potentiation SBA assays, Gc-antibody-serum mixtures were incubated with 80μL of the indicated final antimicrobial concentrations (in PBS for antibiotics or sterile water for lysozyme), and incubated at 37°C, 5% CO_2_ for 2 hr. Samples were then serially diluted and plated on GCB agar for CFU enumeration after overnight culture at 37°C, 5% CO_2_. Where indicated, OMCI (20μg/mL final concentration) or equal volume of TBS buffer was incubated with serum for 30min at 4°C prior to adding Gc.

### Potentiation indexes

For each antimicrobial concentration and serum percentage, CFU enumerated from serum alone was divided by the CFU from serum with the antimicrobial. Similarly, CFU enumerated from HI serum alone was divided by CFU from HI serum with the antimicrobial. The potentiation index is the ratio of the effect of antibiotic on active serum vs. HI serum:

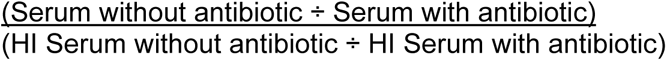

A potentiation index >1.0 indicates a greater-than-additive effect of combining active serum and antimicrobial, while a potentiation index ≤ 1.0 indicates no enhanced effect.

### Complement deposition by imaging flow cytometry

Bacteria from GCB plates were inoculated into HBSS-BSA to an OD_550_ nm of 0.25 and mixed 1:1 with IgM for 30min at 37°C, 5% CO_2_. Buffer or serum were then added (final serum concentration of 2% for C3, or 50% for C7 and C9) and incubated for 2hr more. Bacteria were washed three times with PBS (for C3 and C9) or HBSS-BSA (for C7). For C7, AlexaFluor 488-conjugated (AF488) anti-IgG was then added for 30min at 4°C in the dark, then washed into PBS. Bacteria were counterstained with Tag-it Violet (TIV; BioLegend) for 15min at 37°C with 5% CO_2_, washed into buffer, and fixed with 1% paraformaldehyde overnight. Samples were assayed using the Imagestream^X^ Mk II with INSPIRE software (Luminex) within 72hr. FITC and AF488 were detected with excitation at 488nm and 480–560nm emission; PE with 561-nm laser excitation and 560–561nm emission; and TIV with excitation at 405 nm and 420–505nm emission. Single-color fluorescence samples were collected without brightfield or scatter to create compensation matrices for each experiment and aid in gate-setting. All events (10,000 per sample) were collected on focused singlet cell events and micrographically verified as described (59). Results are presented as the fluorescence index, defined as the median fluorescence intensity multiplied by the percentage of positive-gated bacteria.

### Fluorometric measurements of bacterial membrane integrity

Gc was inoculated from plates into HBSS-BSA to an OD_550_ nm of 0.1. IgM (90μL) was added to 90μL of Gc for 30min at 37°C, 5% CO_2_. For NPN, Gc was then incubated with 50% final concentration IgG/M-depleted serum for 15min (lot #15443); for SYTOX Green, Gc was incubated with 2% IgG/M-depleted serum for 30min (lot #28341). Bacteria were washed three times with buffer and resuspended in 30μM NPN (Sigma-Aldrich, Cat. #104043) (57, 64) or 10μM SYTOX Green nucleic acid stain (Sytox Green; Invitrogen, Cat. #S7020) (19, 55), respectively. Bacteria were resuspended, transferred to black flat-bottom 96-well plates in 100μL technical duplicates, and assayed immediately. NPN measurements were collected on a BioTek Synergy2 plate reader with Gen5 software using 360nm excitation and 420-480nm emission. Sytox Green was measured every 2-4 min over 120 min at 37°C on a PerkinElmer Victor^3^ 1420 Multilabel Counter with associated software, using 490nm excitation and 535nm emission filters. Each experiment included buffer-alone and NPN/Sytox Green without bacteria controls (i.e. blanks), the values of which were subtracted from experimental conditions.

### Minimum inhibitory concentrations (MICs)

100μL of IgG/M-depleted human serum, diluted to 10% in GCBL with Kellogg’s supplement I (121) and 1.25μM Fe(NO_3_)_3_ (GCBL+Supp), was added to each well in one row of a round-bottom 96-well plate. Wells in the next row were filled with 100μL GCBL+Supp (0% serum). 100μL of antimicrobials (4x final concentration) were added to the second column of each row, leaving the first column as no-antimicrobial control. Antimicrobials were then serially diluted 2-fold across the remaining wells in each row. To the no-antimicrobial wells, 100μL of GCBL+Supp was added and mixed thoroughly, and 100μL was removed and discarded. Gc was inoculated into GCBL+Supp to a final OD_550_ nm of 0.07, diluted 10-fold (∼5e6 CFU/mL), and 100μL added to each well and mixed thoroughly. After incubation for 16hr at 37°C, 5% CO_2_. wells were gently resuspended and assessed visually for gonococcal growth, from which MICs were determined (131).

### Statistics, analyses, and data availability

Results are depicted as mean ± standard error for ≥ 3 independent replicates. Statistics were calculated and data were graphed using GraphPad Prism. Data were assumed to be parametric, and statistical tests were 2-sided where applicable. Data and statistics for flow cytometry were obtained using IDEAS 6.2 software (Amnis). Raw data are available from the authors upon request.

## Acknowledgements and Sources of Funding

We thank past and present members of the Criss Lab for advice and insights. We thank Keena Thomas (UVA) and the UVA Flow Cytometry Core Facility for assistance with imaging flow cytometry data acquisition and analysis. Sanjay Ram (University of Massachusetts), Anna Blom and Frida Mohlin (Lund University), Hank Seifert (Northwestern University), and Dan Gioeli (UVA) generously provided bacterial strains, reagents, and access to equipment. We are grateful to Ron Taylor (UVA) for perspective and insight on this project. This work was supported by NIH R01 AI097312, U19 AI144180, and the UVA Harrison Distinguished Teaching Professorship (AKC). ERL was supported by NIH F30 AI179038, T32 AI007046, and T32 GM007267.

## Authorship Contribution Statement

<u>ERL:</u> Conceptualization, Methodology, Analysis, Investigation, Writing – Original Draft & Editing, Validation, Visualization, Funding Acquisition. <u>AKC:</u> Conceptualization, Methodology, Analysis, Writing – Original Draft & Editing, Funding Acquisition, Project Administration, Supervision.

## Supplemental Figure Legends

**Supplemental Figure 1. Vancomycin activity is potentiated by serum in the MS11 strain of Gc.** MS11 Gc was preincubated with anti-Gc IgM, followed by incubation with 2% IgG/M-depleted human serum with or without prior heat inactivation (HI), and subsequent incubation with 4μg/mL vancomycin or vehicle alone. Error bars are standard error of the mean of enumerated CFU. Significance was determined by 1-way ANOVA with Tukey’s multiple comparisons on Log_10_-transformed data. * = p<0.05, **** = p<0.0001. Dotted line represents CFU limit of detection.

**Supplemental Figure 2. For multiple strains of Gc, serum decreases the minimum inhibitory concentrations of nisin and ceftriaxone.** The indicated Gc strains were assayed via 16-hour minimum inhibitory concentration (MIC) broth microdilution assays over a range of nisin or ceftriaxone concentrations in GCBL alone, or supplemented with 2.5% human serum.

